# CRISPR/Cas9-mediated identification of human macrophage SphK1 as druggable target for development of anti-leishmanial chemotherapeutics

**DOI:** 10.1101/2025.08.18.670766

**Authors:** Evanka Madan, Jhalak Singhal, Aashima Gupta, Soumyadeep Mukherjee, Waseem Dar, Neha Jha, Pragya Gupta, Soumya Pati, Sivaprakash Ramalingam, Shailja Singh

## Abstract

Sphingosine-1-phosphate (S1P) is a bioactive lipid that regulates apoptosis, autophagy, inflammation, and intracellular pathogen survival. While S1P signaling has been implicated in *Leishmania donovani* infection, the specific roles of its biosynthetic enzymes Sphingosine kinases SphK1 and SphK2 in host macrophage remains poorly defined. Here, we delineate the role of SphK1 in modulating host-pathogen interactions using CRISPR/Cas9-mediated knockout and pharmacological inhibition in THP-1 macrophages.

We generated and validated CRISPR constructs targeting SphK1 (promoter, SBS, exon 6) and SphK2 (exon 3). SphK1 Knockout was confirmed at transcript and protein levels, accompanied by a marked reduction in SphK1 enzymatic activity and S1P levels. Functionally, SphK1 knockout macrophages exhibited decreased intracellular *L. donovani* burden, elevated TNF-α, and reduced IL-10, and increased autophagic and apoptotic markers, suggesting a pro-inflammatory, cell-death-prone state.

Pharmacological inhibition using the selective SphK1 inhibitor, PF-543 recapitulated these findings, showing reduced phosphorylated SphK1, enhanced p38MAPK activation, and augmented autophagy and apoptosis. Conversely, the SphK2 inhibitor ABC294640 had minimal effect, reinforcing the predominant role of SphK1.

Together, our study identifies SphK1 as a critical host factor that facilitates *L. donovani* survival by modulating lysosomal stress, immune evasion, and cell fate pathways. Targeting SphK1-S1P signalling may offer a novel therapeutic approach for visceral leishmaniasis.

## Introduction

Leishmaniasis is a neglected tropical disease caused by intracellular protozoa parasite *Leishmania* which primarily infects and resides within host macrophages. The parasite manipulates host immune responses and rewires metabolic pathways, particularly those involving lipid metabolism pathways, to create a permissive intracellular environment (1,2).

One such critical lipid signalling pathway is Sphingosine-1-phosphate (SIP) signalling axis. The cascade begins with the generation of S1P from sphingosine by the action of SphK. Afterward, S1P translocates to the outer membrane and binds to its receptors, namely S1PR1-5, which triggers the small G-proteins associated with them. These cascades regulate numerous cellular processes, including proliferation, differentiation, survival, and immune cell trafficking (3,4). Disruption of S1P signalling has been linked to a wide range of pathological conditions, including inflammation, fibrosis, asthma, ischemia-reperfusion injury, and certain cancers (5,6). Notably, studies have indicated that inhalation of SphK1 inhibitors attenuates airway inflammation in murine asthma models, underscoring its therapeutic potential (7,8).

Among the two isoforms, SphK1 is particularly responsive to extracellular stimuli and has emerged as a more viable therapeutic target than the constitutively active SphK2 (9). SphK1 not only regulates host S1P levels but is also exploited by *Leishmania* for detoxification, stress resistance, and virulence, making it a pleiotropic enzyme essential to both host and parasite biology (10,11). Thus, modulating SphK1 activity may be an effective strategy to limit *Leishmania* infection.

Based on the fact that SphK1 is essential for the intracellular survival of *Leishmania donovani* within human macrophages, we conjectured that genetic disruption of this enzyme will impair parasite survival by promoting autophagy, apoptosis, and pro-inflammatory responses, thereby establishing SphK1 as a viable host-directed therapeutic target for anti-leishmanial chemotherapeutics.

To test our hypothesis, we used CRISPR/Cas9 gene editing technology to selectively disrupt SphK1 and SphK2 genes in THP-1 macrophages using plasmid PX459-turboGFP. sgRNAs were designed for targeting key regions of SphK1 (promoter, substrate binding site, exon 6) and SphK2 (exon 3) and validated. Knockout effects were evaluated via autophagy and apoptosis markers (LC3, Beclin1, Cytochrome C, Caspase 3/9, Atg5) in wild-type, vector-only, and sgRNA-transfected cells. Additionally, SphK1 inhibitors: PF-543, ABC294640, were tested on *Leishmania*-infected THP-1 macrophages to study their effects on intracellular parasite burden and host response. Given the limited efficacy of current anti-leishmanial therapies, increasing resistance, and lack of new therapeutic agents, targeting host SphK1 offers a novel, host-directed strategy. Notably, because SphK1 is a host-encoded enzyme, the risk of parasite-driven resistance is minimized. Moreover, published evidence and our observations suggest that SphK1 deletion does not adversely affect macrophage viability, making it a promising target for therapeutic intervention (10,11).

To our knowledge, this study is the first to comprehensively evaluate the role of SphK1 inhibition and genetic deletion in regulating S1P signalling, lysosomal stress, and parasite survival within human macrophages infected with *Leishmania donovani*. By integrating small-molecule inhibition with CRISPR/Cas9-mediated gene editing, our findings offer critical mechanistic insights and potential translational implications for the development of host-targeted anti-leishmanial therapies.

## Results

### 1. Validation of SphK1 and SphK2 CRISPR constructs via PCR, Sanger Sequencing, and GFP expression

To genetically ablate SphK1 and SphK2 expression in THP-1 macrophages, we designed multiple CRISPR/Cas9 constructs targeting key regulatory and coding regions of the respective genes. Specifically, guide RNAs (sgRNAs) were designed to target the substrate-binding site, promoter region, and exon 6 of the SphK1 gene, and exon 3 of the SphK2 gene. Each sgRNA was cloned into the PX459-TurboGFP expression vector, which allows simultaneous Cas9 expression and GFP-based tracking of transfection efficiency **(Fig S1)** in line with previous studies (12, 13).

Following transformation into competent *E. coli*, six colonies per construct were randomly selected for colony PCR using a plasmid-specific U6 forward primer and a guide RNA-specific reverse primer to verify the presence of the desired insert. PCR amplification yielded ∼222 bp fragments, confirming successful cloning of the sgRNA inserts **(Fig S2A–D (i)).**

Subsequently, plasmid DNA from positive colonies was extracted and subjected to Sanger sequencing to ensure the fidelity and correct orientation of the cloned guide sequences. The sequencing data confirmed accurate integration of sgRNAs with intact PAM sequences and no frame-shift mutations or mismatches (14), thereby validating the constructs for downstream applications **(Fig S2A–D (ii))**.

To assess transfection efficiency and expression of the CRISPR constructs in mammalian cells, each sgRNA construct was transfected into THP-1 macrophages. GFP fluorescence, driven by the TurboGFP reporter in the PX459 vector backbone, was used to identify successfully transfected cells via fluorescence microscopy. Robust GFP signal was observed 48–72 h post-transfection in multiple fields of view, indicating efficient delivery and expression of the sgRNA-Cas9 machinery in THP-1 cells **(Fig S2A-D (iii)).**

These results collectively confirm successful molecular cloning, sequence fidelity, and functional transfection of SphK1 and SphK2 CRISPR constructs into host macrophages, establishing a validated system for gene editing and subsequent functional studies.

### 2. Targeted disruption of SphK1 in macrophages reduces mRNA expression, protein levels, and S1P formation

Guide RNAs designs and their genomic sequences targeting the Promoter region, the Substrate Binding Site and Exon-6 of the hSphK1 gene (4652 bp) is shown **(Fig 1A).** To evaluate the efficacy of SphK1-targeting sgRNAs, initial experiments were conducted in THP-1-derived macrophages. *In vitro* transcription of the sgRNAs targeting the SphK1 gene was performed as shown in **Fig S3 (i).** The T7 endonuclease I assay demonstrated robust cleavage activity in DNA templates corresponding to the promoter and substrate-binding site (SBS) regions of the SphK1 gene when incubated with recombinant Cas9 and the respective sgRNAs, indicating high editing efficiency **(Fig S3 (ii)).**

**Fig 1.**
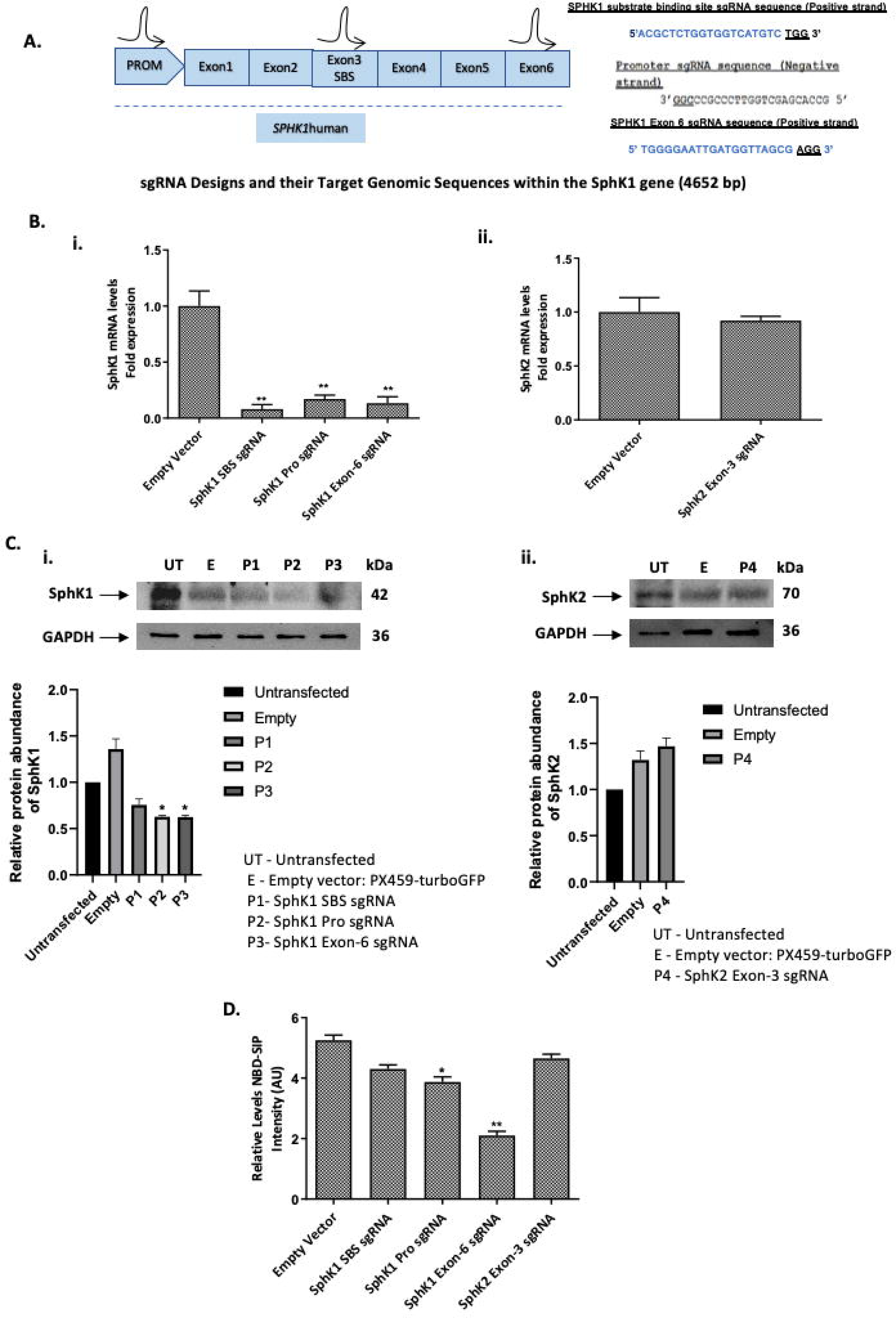
Characterization of SphK1 and SphK2 Knockout in THP-1 Macrophages. **(A)** Schematic representation of sgRNA designs and their respective target genomic loci within the human SphK1 gene (4652 bp). **(B)** qRT-PCR analysis of *SphK1* and *SphK2* mRNA levels following sgRNA-mediated knockout in THP-1 macrophages. THP-1 macrophages were cultured in 25 cm^2^ flasks followed by transient transfection of 10µg of sgRNAs targeting *SphK1* (Promoter, SBS, Exon-6) and *SphK2* (Exon-3) for 72h. Total RNA was extracted from transfectants using Trizol, and the resulting cDNA was subjected to real-time PCR analysis using primers specific for human *SphK1* and *SphK2*. qRT-PCR analysis showing relative mRNA levels of *SphK1* and *SphK2* post-transfection. The results are expressed as fold-change relative to empty vector control. *RNU6A* was used as a housekeeping gene. Values are mean ± S.D (n = 3). The results are representative of three independent experiments performed in triplicates. (**C)** Western blot analysis of SphK1 and SphK2 protein levels post-transfection. Total protein lysates were subjected to SDS-PAGE and immunoblotted with anti-SphK1 and anti-SphK2 antibodies. GAPDH was used as a loading control. The intensity of the bands was quantified by densitometry using AlphaEase FC Imager software. The densitometric analysis shows the fold change in expression relative to control. Values are mean ± S.D. (n = 3). (**D)** Assessment of NBD-S1P production in SphK1-or SphK2-knockout THP-1 macrophages. Cells were transfected with sgRNA constructs for 72h, then incubated with 10µM NBD-sphingosine for 45 min at 37 °C. Macrophages phosphorylate NBD-sphingosine to NBD-S1P via SphK1 activity. Cells were processed in BSA-containing buffer for uptake and quantification. Bar graph depicts ELISA-based quantification of S1P. SphK1 knockout significantly reduced S1P production, with no notable change in SphK2 knockout cells. Results represent three independent experiments. (UT: Untransfected THP-1 cells, E: Empty vector PX459-turboGFP, P1-SphK1-SBS sgRNA, P2-SphK1-Pro sgRNA, P3-SphK1-Exon-6 sgRNA, P4-SphK2-Exon-3 sgRNA).

Gene knockout efficiency was further validated at the transcript level by quantitative RT-PCR. using primers specific for *SphK1 and SphK2* in SphK knockout THP-1 cells. THP-1 cells transfected with sgRNAs targeting the promoter, SBS, and exon 6 regions of SphK1 exhibited a marked reduction (∼90%) in SphK1 mRNA expression compared to controls (Empty Vector). Notably, SphK2 mRNA levels remained unaffected in cells transfected with sgRNAs targeting exon 3, confirming the specificity of the sgRNAs **(Fig 1B).**

Western blot analysis corroborated the transcript-level findings. Western blot analysis using cell lysates derived from untransfected THP-1 **(Fig 1C, lane 1, UT),** Empty vector: PX459-turboGFP **(Fig 1C, lane 2, E),** THP-1 cells transfected with SphK1 SBS sgRNA **(Fig 1C, lane 3, P1)**, THP-1 cells transfected with SphK1 Pro sgRNA **(Fig 1C, lane 4, P2),** THP-1 cells transfected with SphK1 Exon-6 sgRNA **(Fig 1C, lane 5, P3)** was performed. Immunoblotting was done using anti-SphK1 and SphK2 antibodies. Cells transfected with sgRNAs targeting the promoter, SBS, and exon 6 regions showed substantially decreased SphK1 protein expression relative to control cells. In contrast, no changes were observed in SphK2 protein expression in cells transfected with exon 3-targeting sgRNAs **(P4) (Fig 1C)** in line with previously reported findings (15, 16).

To assess the functional consequences of SphK1 gene disruption, a fluorimetry-based SphK activity assay was performed using NBD-sphingosine as the substrate. A significant reduction in NBD-S1P formation was observed across all SphK1 sgRNA-transfected groups, with the most pronounced inhibition detected in cells targeting exon 6 **(Fig 1D)**. Correspondingly, fluorescence intensity was markedly decreased in promoter, SBS, and exon-6 targeted SphK1 sgRNA transfectants relative to the empty vector control, indicating effective suppression of SphK1 enzymatic activity.

### 3. Reduced SphK1 fluorescence reflects functional loss of S1P synthesis following gene knockout in macrophages

In line with the above findings, fluorescence intensity measurements from the SphK activity assay demonstrated a significant reduction in enzymatic activity in THP-1 cells transfected with sgRNAs targeting the promoter, substrate-binding site (SBS), and exon 6 regions of the SphK1 gene, compared to cells transfected with the empty vector control **(Fig 2A).** In contrast, no appreciable changes in fluorescence were observed in THP-1 cells transfected with sgRNAs targeting the exon 3 region of the SphK2 gene **(Fig 2B),** confirming the specificity of SphK1 targeting.

**Fig 2.**
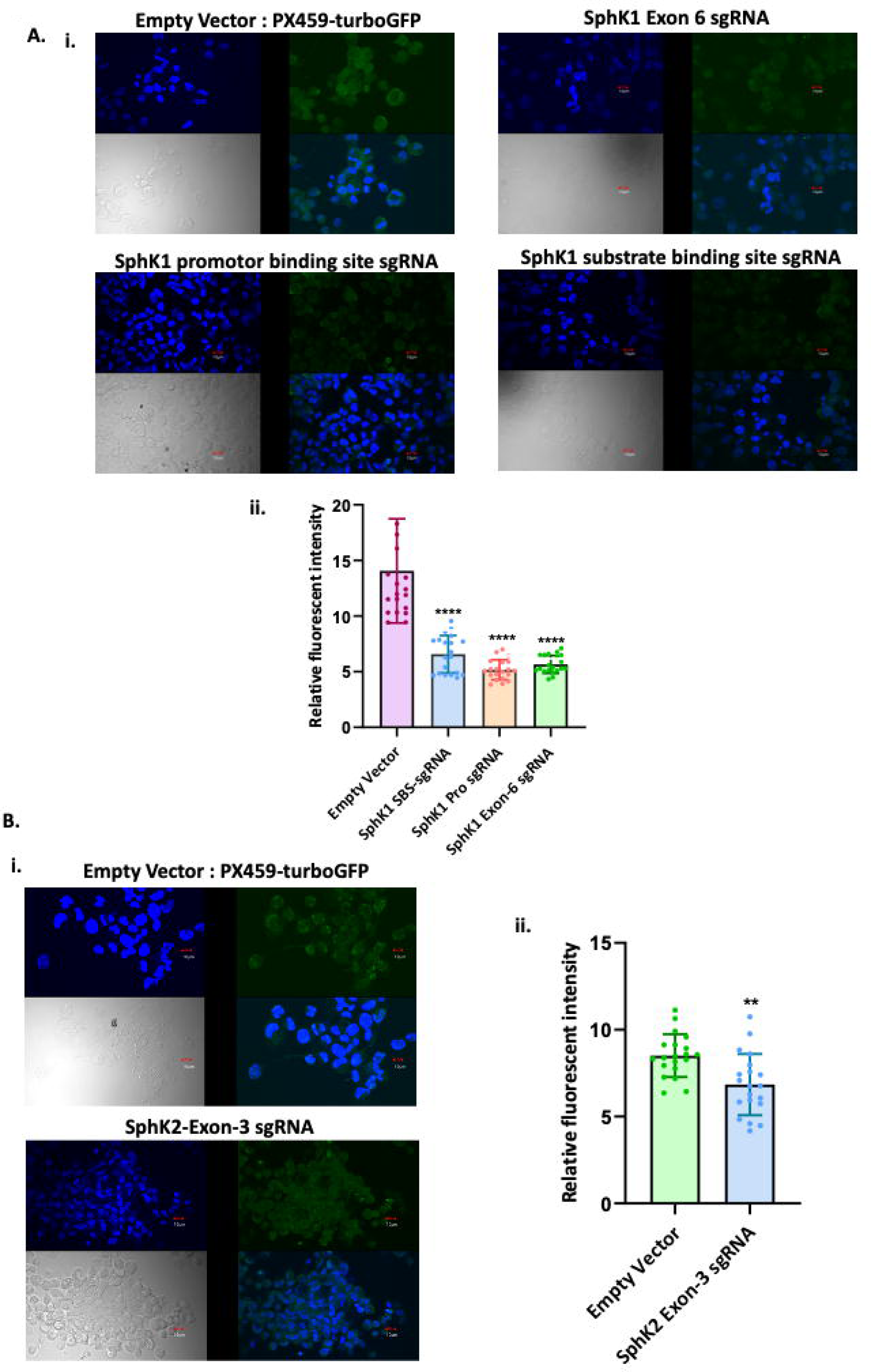
Confocal Imaging of SphK1 and SphK2 expression and localization in SphK knockout THP-1 macrophages. **(A)** Representative confocal micrographs showing SphK1 expression and its subcellular localization in THP-1 macrophages transfected with empty vector: PX459-turboGFP, SphK1 Exon 6 sgRNA, promotor binding site sgRNA, substrate binding site sgRNA constructs. (**B)** Confocal micrographs displaying SphK2 expression in THP-1 macrophages transfected with either the empty vector: PX459-turboGFP or SphK2 Exon 3 sgRNA. Mean fluorescence intensity (MFI) of individual transfected macrophages was quantified and plotted relative to the empty vector-transfected controls. Data are presented as mean ± SD from three independent experiments (n = 3). Statistical significance was determined using appropriate tests; p ≤ 0.01.

The observed decrease in fluorescence intensity reflects a reduction in the conversion of NBD-sphingosine to NBD-S1P, indicating impaired SphK1 enzymatic activity. Among all targeted sites, exon 6-targeted cells **(P3)** exhibited the most marked suppression of fluorescence, consistent with the most effective inhibition of SphK1 catalytic function. This aligns with previous work implicating exon-6 as crucial for catalytic activity (17). These findings collectively confirm that CRISPR/Cas9-mediated editing at the promoter, SBS, and exon 6 loci of the SphK1 gene results in a substantial loss of both gene expression and enzymatic function, thereby validating the functional impact of gene disruption at these specific sites.

### 4. SphK1 knockout reduces intracellular *Leishmania* burden and triggers pro-inflammatory, autophagic and apoptotic responses in THP-1 macrophages

To assess the role of SphK1 in controlling *Leishmania donovani* infection, THP-1 macrophages were transfected with sgRNAs targeting the promoter, substrate-binding site (SBS), and exon 6 regions of the SphK1 gene before infection at a multiplicity of infection (MOI) of 20:1. To determine, the intracellular parasite burdens (mean number of amastigotes per macrophage) were microscopically assessed using Giemsa staining. Untransfected, infected THP-1 cells served as positive controls and showed an infection rate of approximately 80%. In contrast, SphK1-knockout cells; THP-1 cells transfected with sgRNAs targeting the promoter, substrate-binding site (SBS), and exon 6 regions of the SphK1 gene, compared to cells transfected with the empty vector control exhibited significantly reduced infection rates, with ∼30–40% of cells harbouring amastigotes **(Fig 3A)** consistent with the role of host lipid signalling in parasite survival (18, 19).

**Fig 3.**
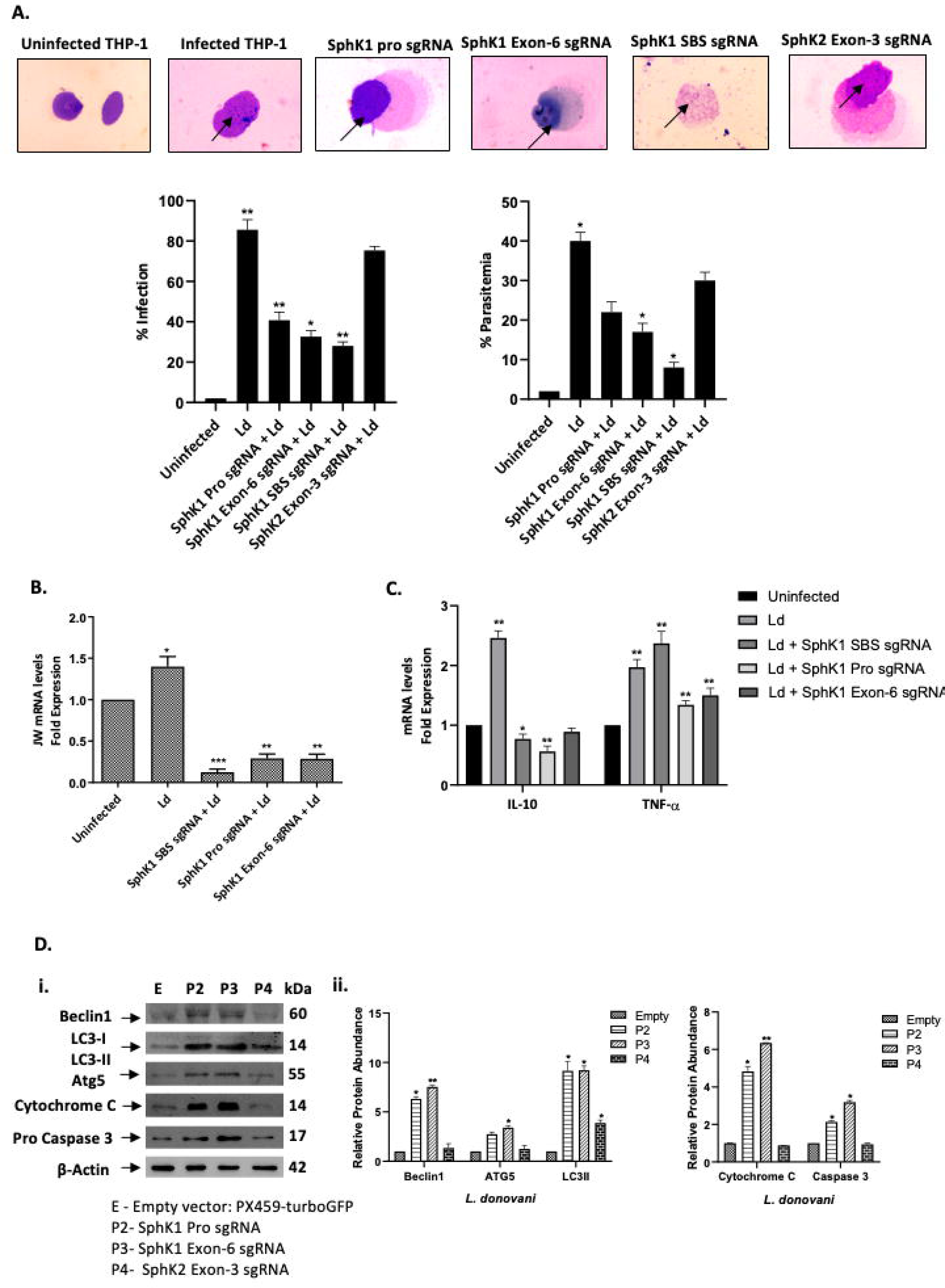
CRISPR-Cas9-mediated knockout of SphK1 reduces parasite burden, induces a pro-inflammatory cytokine profile, and enhances autophagy and apoptosis in *Leishmania*-infected THP-1 macrophages (A, **B)** SphK1 knockout reduces *L. donovani* infection in THP-1 macrophages. THP-1 cells grown in RPMI medium were transfected with sgRNAs targeting SphK1 (substrate-binding site, promoter, exon 6) or SphK2 (exon 3), or with the empty vector (PX459-turboGFP) for 24h, followed by infection with *L. donovani* promastigotes (MOI = 20:1) for 48h. (**A)** 48h post-infection, cells were Giemsa stained and examined microscopically for amastigotes. Virulence capacity was determined by calculating infectivity (left panel) and parasitaemia (right panel). THP-1 cells transfected with empty vector: PX459-turboGFP and infected with WT-*L. donovani* were used as control. The results are representative of three independent experiments. The results signify mean ± S.D with n = 3, *P < 0.05 statistical difference from the infected wild-type control. (**B)** Parasite burden was quantified via qRT-PCR targeting the parasite-specific *JW* gene. Total RNA was enriched using TRIzol and the resulting cDNA was subjected to real-time PCR analysis using primers specific for infectivity; *JW* gene in transfected, infected and uninfected macrophages. *RNU6A* was used as a housekeeping gene. The results are expressed as fold-change relative to uninfected THP-1 cells. Statistical significance was quantified using the unpaired t-test. The data is a representation of mean ± SD from three independent experiments *, p< 0.05; ***, p < 0.001. **(C)** SphK1 knockout promotes a pro-inflammatory cytokine response. At 48h post-infection, total RNA was extracted and analyzed for *IL-10* and *TNF-*α transcript levels via qRT-PCR. Results are presented as fold-change relative to uninfected, untransfected controls. Data were analyzed using the 2^–ΔΔCT method. RNU6A was used as an internal control. Mean ± SD (n = 3), representative of three independent experiments. *p < 0.05 vs infected PX459 control. **(D)** SphK1 knockout enhances autophagy and apoptosis pathways in infected macrophages. **(i)** Western blot analysis of autophagy markers (Beclin-1, LC3-I/II, Atg5) and apoptosis markers (Cytochrome c, Pro-Caspase 3) in THP-1 macrophages harvested 72h post-transfection and 48h post-infection. **(ii)** Densitometric quantification of band intensities was performed using ImageJ and plotted with GraphPad Prism 8. Results represent mean ± SD from three independent experiments. *p < 0.05; ***p < 0.001 (unpaired t-test).

To confirm these observations, qRT-PCR targeting the parasite-specific kinetoplast minicircle gene (JW) was performed. SphK1 knockout THP-1 cells exhibited a significant decrease in parasitic RNA compared to untransfected, infected controls **(Fig 3B),** supporting the conclusion that SphK1 is crucial for parasite sustenance within the host intracellular environment.

Quantitative analysis of intracellular parasite burden at 48h post-infection, using both Giemsa staining and RT-PCR, revealed a ∼35% reduction in the number of amastigotes per macrophage in cells transfected with SphK1-targeting sgRNAs **(Fig 3A,B).** These findings indicate that SphK1 depletion hampers *L. donovani* survival within host macrophages, likely due to impaired intracellular growth and persistence of the parasite.

The impact of SphK1 disruption on the inflammatory response was evaluated by analysing cytokine gene expression using primers specific for *IL-10 and TNF-*α in SphK knockout THP-1 cells. Transfection with SphK1-targeting sgRNAs resulted in a ∼35% reduction in the anti-inflammatory cytokine, IL-10 and a concurrent increase in TNF-α transcript levels (∼ 2.5 folds) **(Fig 3C).** This shift toward a pro-inflammatory cytokine profile (20, 21) suggests that SphK1 knockout may enhance the host’s immune response against *L. donovani*.

To explore whether host cell death or survival pathways were involved in SphK1-mediated effects on parasitaemia, we examined markers of autophagy and apoptosis in infected THP-1 cells. Western blot analysis using cell lysates derived from Empty vector: PX459-turboGFP **(Fig 3D, lane 1, E),** THP-1 cells transfected with SphK1 Pro sgRNA **(Fig 1C, lane 2, P2),** THP-1 cells transfected with SphK1 Exon-6 sgRNA **(Fig 1C, lane 3, P3)** and THP-1 cells transfected with SphK2 Exon-3 sgRNA **(Fig 1C, lane 4, P4)** was performed. Immunoblotting was done using anti-Atg5, Beclin1, LC3-I/II, cytochrome c, pro-caspase 3, and caspase 9 antibodies. Western blot analysis revealed upregulation of autophagy-associated proteins Atg5 (∼3 folds), Beclin1 (∼8 folds), and LC3-I/II (∼10 folds), along with increased expression of apoptotic markers including cytochrome c (∼6 folds), pro-caspase 3 (∼4 folds) in SphK1-knockout macrophages **(P2 and P3) (Fig 3D)** consistent with known roles of SphK1 in regulating autophagy-apoptosis crosstalk (22, 23).

In contrast, no significant changes were observed in cells transfected with sgRNAs targeting the exon 3 region of the SphK2 gene **(P4),** reinforcing the specificity of SphK1 in mediating these effects. These results suggest that loss of SphK1 enhances autophagy and apoptosis pathways in host macrophages, contributing to reduced intracellular parasite survival.

### 5. Pharmacological inhibition of host SphK1 enhances autophagy and apoptosis in *Leishmania*-infected macrophages

The last set of experiments was aimed at identifying the role, if any, of *L. donovani* infection on SphK1 in infected macrophages. THP-1 cells were infected with *L. donovani* in RPMI medium and harvested for Western blot analysis. At 48h post-infection, total protein was extracted from infected macrophages, and subjected to Western blot, using SphK (total and phospho) specific antibody as reported in the methods section. SphK1 expression increased (∼2 folds) (**Fig 4A)** following *L. donovani* infection, consistent with previous reports that parasitic infections manipulate host lipid kinases for survival (24, 25). However, pharmacological inhibition with PF-543 (17µM), a selective SphK1 inhibitor, resulted in a significant decrease (p ≤ 0.01) in both total and phosphorylated SphK1 levels at 48h post-infection, in agreement with its specificity (26, 27). In contrast, treatment with ABC294640 (70µM), a SphK2 inhibitor, did not alter SphK1 expression **(Fig 4A (i, ii))**.

**Fig 4.**
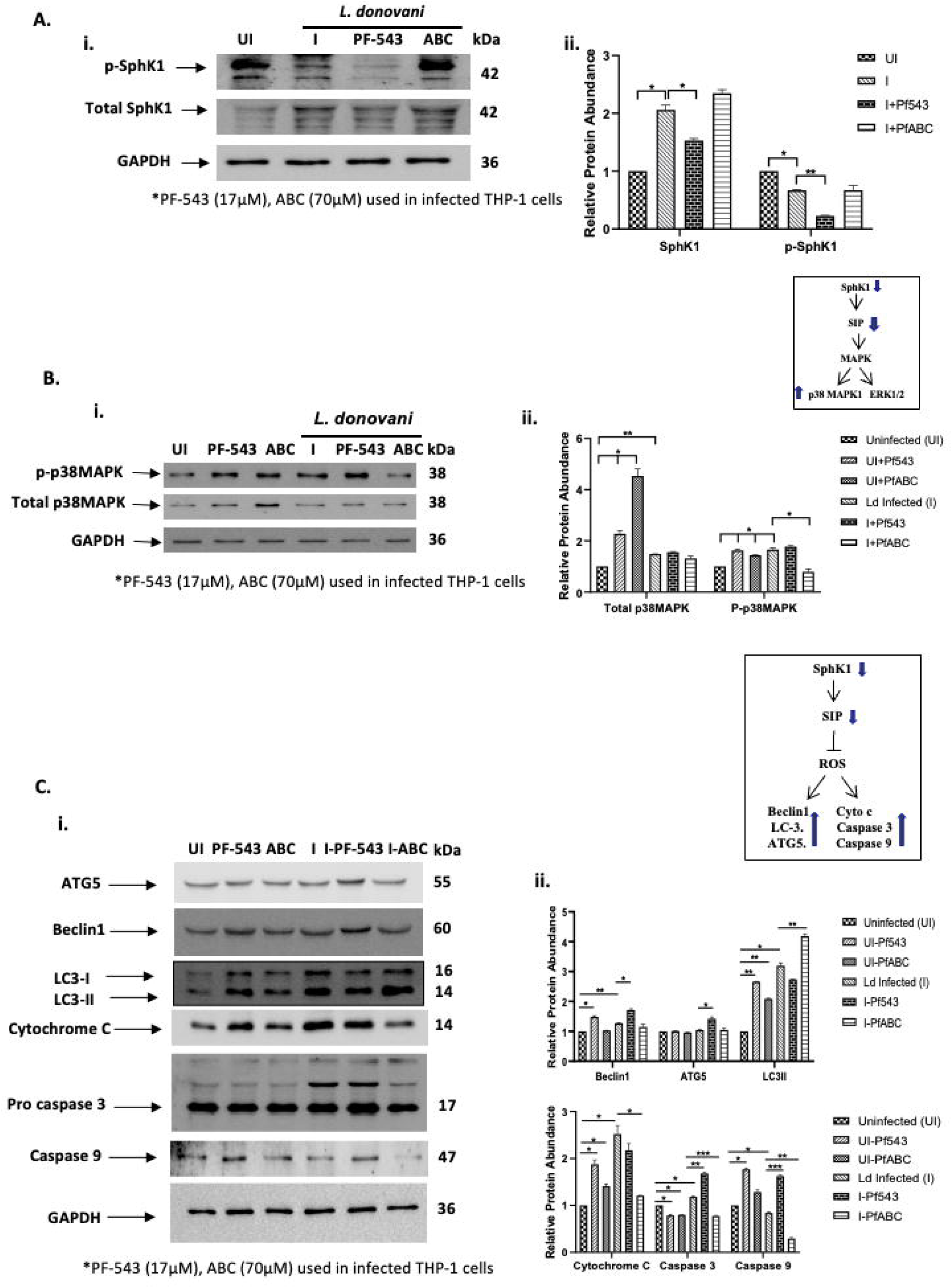
Pharmacological inhibition of host SphK1 attenuates SphK1 signalling and enhances MAPK activation, autophagy, and apoptosis in *Leishmania-*infected macrophages. (A) SphK1 expression in response to infection and inhibition. THP-1 macrophages were cultured in six-well plates in the presence or absence of *L. donovani* infection (MOI = 20:1) for 6h. Infected THP-1 were washed to remove non-internalized parasites and treated with PF-543 (17µM) or ABC294640 (70µM) for 48h. **(i)** Western blot analysis showing total and phosphorylated SphK1 (p-SphK1) in uninfected, infected, and inhibitor (PF-543 and ABC294640)-treated cells. **(ii).** Quantitative densitometric analysis showing fold changes in SphK1 levels during *L. donovani* infection, 48h post treatment with PF-543 or ABC294640. (B) Modulation of MAPK Signalling. **(i)** Western Blot analysis of total and phosphorylated p38MAPK in uninfected, infected, and inhibitor-treated THP-1 macrophages **(ii)** Quantitative analysis showing fold changes in p38MAPK activation in response to PF-543 and ABC294640. **(C)** Induction of autophagy and apoptosis. **(i)** Expression of autophagic markers (Beclin-1, Atg5, LC3-I/II) and apoptotic markers (Cytochrome c, Pro-Caspase 3, Caspase 9) was evaluated by Western blot in uninfected, infected, and inhibitor-treated THP-1 cells at 48h post-infection. **(ii)** Band intensities were quantified using ImageJ, and statistical analysis was performed using GraphPad Prism 8. Data are expressed as mean ± SD from three independent experiments*. *p < 0.05; \*\**p < 0.001.

As reported in literature, since MAPK pathways influence parasite survival, we next analysed p38MAPK activation. At 48h post-infection, total protein was extracted from infected macrophages, and was subjected to Western blot, using p38MAPK (total and phospho) specific antibody. PF-543 treatment enhanced p38MAPK phosphorylation (p ≤ 0.05) in infected THP-1 cells, whereas ABC294640 reduced it **(Fig 4B (i,ii)**), highlighting SphK1’s link to MAPK signaling pathways (28).

To further investigate the role of SphK1 in regulating the expression of autophagy and apoptosis, we examined the status of autophagic, apoptotic markers in intracellular *L. donovani* derived from PF-543 and ABC294640 treated infected THP-1 cells cultured in RPMI medium. Western blot analysis using cell lysates derived from uninfected THP-1 **(Fig 4C, lane 1),** *L. donovani-*infected THP-1 **(Fig 4C, lane 4),** uninfected and PF-543 or ABC294640 treated THP-1 **(Fig 4C, lane 2 or 3)**, *L. donovani*-infected and PF-543 or ABC294640 treated THP-1 **(Fig 4C, lane 5 or 6)** was performed. Immunoblotting was done using anti-ATG5, anti-Beclin-1, anti-LC3I/II, anti-Cytochrome C, anti-Pro-Caspase 3 and anti-Caspase 9 antibodies. **Fig 4C** shows that infection alone increased Beclin-1, LC3-I/II, Cytochrome C, and Caspase 3 levels (∼2 folds, p ≤ 0.05). PF-543 treatment in infected macrophages further elevated Beclin-1, ATG5, Caspase 3, and Caspase 9 levels (∼3-4 folds), indicating enhanced autophagy and apoptosis, supporting the therapeutic potential of SphK1 inhibition in parasitic infections (29, 30). ABC294640-treated cells showed no significant change compared to untreated infected controls **(Fig 4C (i,ii))**.

## Discussion

This study demonstrates that SphK1 as an essential host factor for intracellular survival of *Leishmania donovani* in human macrophages and establishes a CRISPR–mediated SphK1 knockout in THP-1 cells as a robust platform to probe its function. By designing sgRNAs against both regulatory and coding regions of the SphK1 locus, we achieved efficient gene disruption **(Figs. 1–3, S2),** which yielded a pronounced decrease in SphK1 mRNA and protein expression and virtually abolished cellular sphingosine-1-phosphate production. Functionally, SphK1-deficient THP-1 macrophages exhibited dramatically reduced parasite burden, confirming that *L. donovani* exploits host SphK1 activity for its intracellular survival.

Consistent with previous studies implicating SphK1 in modulating inflammatory responses and immune evasion strategies during infection (31, 32), our data demonstrate that SphK1 knockout results in a significant reduction in intracellular parasite burden (∼35–40%) **(Fig 3A,B),** along with a shift toward a pro-inflammatory phenotype, as evidenced by upregulation of TNF-α and suppression of IL-10 **(Fig 3C).** This is in line with the known immunomodulatory role of S1P, which acts via S1P receptors to dampen macrophage activation and promote Th2 polarization (33, 34).

At the cellular level, SphK1 depletion led to upregulation of autophagic and apoptotic markers, including Beclin1, Atg5, LC3-II, Cytochrome C, and Caspase-3/9 **(Fig 3D)**, suggesting enhanced lysosomal stress responses in infected macrophages. These findings are supported by previous evidence linking SphK1 activity with cell survival pathways and resistance to apoptosis in cancer and infection models (35,36).

Importantly, pharmacological inhibition of SphK1 using PF-543 recapitulated the effects observed with gene knockout, confirming that SphK1 enzymatic activity, rather than expression alone, is essential for parasite survival **(Fig 4A–C).** PF-543-treated cells showed enhanced phosphorylation of p38 MAPK, a stress-responsive kinase linked to pro-inflammatory signalling, further supporting the model of stress-induced immune reactivation upon SphK1 inhibition **(Fig 4B)**. This aligns with the emerging literature suggesting that S1P acts as a brake on p38 and JNK activation in immune cells (37).

Notably, targeting SphK2 did not result in significant changes in parasite burden, S1P levels, or host cell stress responses **(Fig 1–3),** underscoring the isoform-specific role of SphK1 in infection. This finding echoes earlier work showing that SphK2 has a more constitutive role in nuclear regulation and apoptosis, while SphK1 is dynamically regulated and mediates cytoplasmic S1P production during inflammatory stress (38, 39). This isoform-specific role also provides an opportunity for selective targeting, minimizing off-target effects on essential cellular functions.

We propose a working model **(Fig 5)** in which CRISPR-mediated SphK1 knockout or pharmacological inhibition disrupts S1P production, promoting autophagy and intrinsic apoptosis, thereby creating a hostile intracellular environment for *Leishmania*. This shift reduces parasite tolerance and favours immune-mediated clearance. Furthermore, the lack of effect on overall macrophage viability suggests that host-directed therapy targeting SphK1 may offer a low-toxicity strategy to overcome resistance and improve parasite control, without compromising host defence mechanisms.

**Fig 5.**
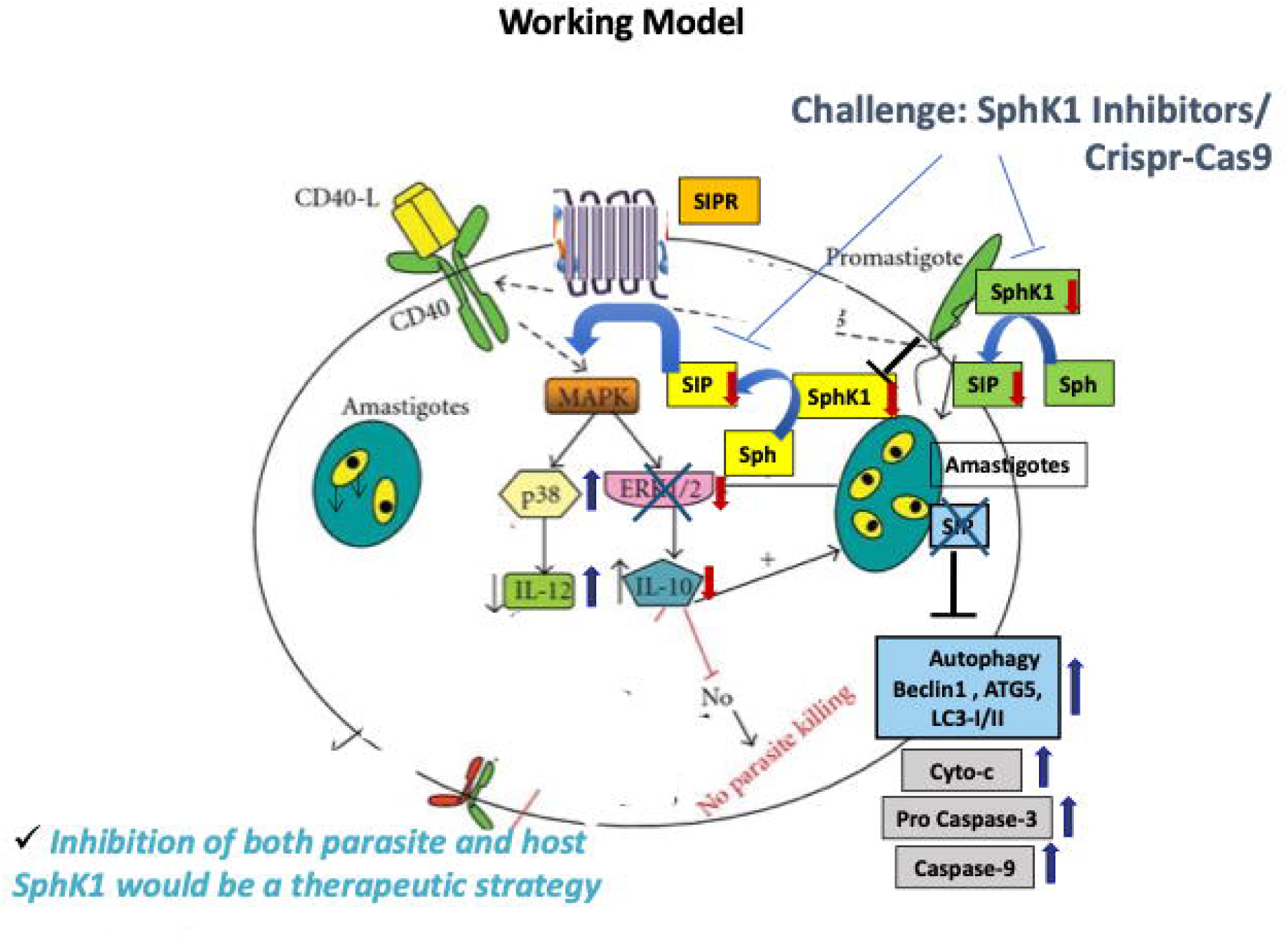
Proposed Model of SphK1-mediated regulation of host cell fate during *Leishmania* infection. Inhibition of SphK1, either pharmacologically using PF-543 or genetically via CRISPR/Cas9 leads to a reduction in phosphorylated SphK1 (p-SphK1) and intracellular S1P levels. This suppression activates the p38 MAPK signalling cascade, resulting in upregulation of IL-12 and downregulation of IL-10, thereby skewing the immune response toward a pro-inflammatory phenotype. Concurrently, enhanced expression of autophagy markers (Atg5, Beclin-1, LC3-I/II) and apoptotic markers (Cytochrome c, Caspase 3, and Caspase 9) facilitates parasite clearance by promoting host cell death pathways.

Taken together, these findings provide compelling evidence that SphK1 is a non-redundant regulator of *Leishmania* persistence and a viable host-targeted therapeutic candidate. Given the emergence of antimonial and miltefosine resistance, host-directed strategies may provide a complementary or alternative approach to traditional antiparasitic therapies (40, 41). Future studies should explore *in vivo* validation of SphK1 inhibition using murine models of leishmaniasis and investigate potential synergies with conventional anti-leishmanial agents.

The broader implications of this study are multi-fold. First, our findings position host SphK1 as a non-redundant regulator of parasite tolerance and macrophage viability. Second, by showing that host-directed SphK1 inhibition does not compromise macrophage survival, this strategy emerges as a low-toxicity alternative with reduced risk for parasite resistance development, as the target is host-encoded. Third, this work bridges the mechanistic gap between sphingolipid metabolism, lysosomal stress, and immune modulation, three axes critical to *Leishmania* pathogenesis.

While these findings highlight important mechanistic insights and therapeutic potential, certain limitations should be considered when interpreting the results. Nevertheless, THP-1 macrophages are a widely used model they may not reflect the complexity of primary human macrophages as well as *in vivo* microenvironment. Secondly, our study primarily focussed on *Leishmania donovani* so these results may not be applicable or can be extrapolated to other *Leishmania* species that have different characteristics and tissue tropisms.

In conclusion, our study provides novel mechanistic insight into the role of host SphK1 in regulating *Leishmania* survival and demonstrates that its genetic deletion or pharmacological inhibition triggers apoptotic and autophagic pathways detrimental to parasite persistence. These findings open new avenues for host-directed therapeutics against leishmaniasis and potentially other intracellular pathogens that exploit host lipid metabolism.

## Materials and Methods

### Materials

THP-1 cells were procured from NCCS, Pune, India. PF-543, ABC294640 and other reagents including RPMI-1640, M199, Fetal Bovine Serum (FBS), penicillin and streptomycin were purchased from Sigma-Aldrich (USA) and Life Technologies. PCR primers were synthesised by Eurofins (USA). Antibodies used include β-actin (Santa Cruz #sc-8432; 1:1,000), Beclin-1 (CST, Danvers, MA, USA; #3495T, 1:1,000), Atg5 (CST #12994T, 1:1,000), and LC3-I/II (CST #4108S, 1:1,000), Caspase-9 Mouse mAb (CST #9508), Caspase-3 (D3R6Y) Rabbit mAb (CST #14220), Cytochrome c (CST #4272), Total/phospho-p38 (Thr180/Tyr182) MAPK (CST # 9211), Total/phospho (Ser-225) Sphingosine kinase 1 (ECM biosciences # SP1641), Anti-Sphingosine Kinase 2 (N-terminal region) Antibody (ECM biosciences # SP4621). Goat anti-mouse horseradish peroxidase (HRP)-conjugated secondary antibody (1:5,000) was purchased from Invitrogen (Carlsbad, CA, USA). Goat anti-rabbit HRP-conjugated secondary antibody (1:2,500) was purchased from ABclonal (Woburn, MA, USA).

### Leishmania cell culture

*Leishmania donovani Bob* strain (*Ld*Bob strain/MHOM/SD/62/1SCL2D) (*WT-L. donovani*) promastigotes, originally obtained from Dr Stephen Beverley (Washington University, St. Louis, MO) were cultured at 26°C in M199 medium (Sigma-Aldrich, USA), supplemented with 10% heat-inactivated fetal bovine serum (FBS; Biowest), 100 units/ml penicillin (Sigma-Aldrich, USA) and 100 µg/ml streptomycin (Sigma-Aldrich, USA). *LdBob* is a genetically modified *Leishmania donovani* strain that is non-virulent in humans but retains the capacity to infect macrophages *in vitro.* It is frequently used as a safe and reliable model organism for studying *Leishmania*–host interactions, parasite biology, and drug screening under standard biosafety level 2 (BSL-2) conditions.

### THP-1 cell culture and infection

THP-1 cells, an acute monocytic leukaemia-derived human cell line (202 TM; American Type Culture Collection, Rockville, MD), were procured from NCCS, Pune in 2022. All cell lines were routinely tested and confirmed to be free of mycoplasma and other contaminants prior to and during the experiments. These procedures ensure that the validity of the experimental results and conclusions remains unaffected. THP-1 cells were maintained in RPMI-1640 medium supplemented with 10% heat-inactivated FBS, 100 units/ml penicillin and 100μg/ml streptomycin at 37°C and 5% CO2. Cells (106 cells/well) were treated with 50 ng/ml phorbol-12-myristate-13-acetate (PMA) (Sigma-Aldrich, USA) for 48h to induce differentiation into macrophage-like-cells before infection. Cells were washed once with phosphate-buffered saline (PBS) and incubated in RPMI medium, 10% heat-inactivated FBS, 100 units/ml penicillin and 100μg/ml streptomycin, before infection. To carry out in vitro infection assays, late stationary phase promastigotes (WT and SphK mutants) were used at a ratio of 20 parasites per macrophage*. Leishmania-*infected macrophages were incubated at 37°C in a 5% CO2-air atmosphere for 4h to allow the establishment of infection and proliferation of intra-macrophage parasites. The cells were then washed five times with PBS to remove non-adherent extracellular parasites. After that, the cells were incubated in RPMI medium at 37°C in a 5% CO2-air atmosphere for 48h. THP-1 cell line widely used as an *in vitro* model to study macrophage biology, innate immune responses, and host–pathogen interactions. It can be differentiated into macrophage-like cells, which closely mimic primary human monocyte-derived macrophages in morphology, surface marker expression, and functional responses, while providing reproducibility and avoiding donor-to-donor variability.

### Generation of SphK1/2 Knockout Macrophages Using CRISPR-Cas9

To disrupt the SphK1 and SphK2 genes in THP-1 cells, sgRNAs were designed to target the human SphK1 gene using the CHOP-CHOP online tool. Literature and knockout-library validated single-guide RNAs (sgRNAs) targeting the SphK1 Substrate Binding Site (https://pmc.ncbi.nlm.nih.gov/articles/PMC7515464/) (42) were selected, and the best sgRNA designs targeting the promoter region and exon-6 of the hSphK1 gene were selected using *in silico* efficiency score calculations **(Table S1).** These sgRNAs were inserted in the PX459-turboGFP vector for co-expression with the *Streptococcus pyogenes* Cas9 protein (A Gift from Sivaprakash Ramalingam Lab at the CSIR-Institute of Genomics and Integrative Biology, New Delhi). The plasmid vector was digested with Bbs1-HF (NEB, Bbs1-HF NEB#R3539). The complementary oligo sequences were hybridized. Double stranded DNA oligos with BbsI restriction overhangs were then ligated into the linearized PX459-turboGFP plasmid featuring complementary BbsI nucleotide overhangs. Plasmid-oligo constructs were then transformed into DH5-alpha competent bacterial strains for amplification on LB-Agar plates supplemented with Ampicillin, 100 μg/mL (Sigma-Aldrich, A9518). Pure bacterial colonies/strains featuring positive constructs were selected by colony PCR screening followed by Sanger sequencing. The human U6 promoter-specific forward primer (5’-GAGGGCCTATTTCCCATGATT-3’) and the oligo insert-specific reverse primers were used for Sanger sequencing confirmation. Expression of positive plasmid constructs in THP-I cells were confirmed by visualizing GFP expression under a fluorescence microscope **(Fig S1**).

### In vitro transcription of the sgRNAs

*In vitro transcription (IVT)* of the sgRNAs were performed using the GeneArt Precision gRNA Synthesis Kit (#A29377; Thermo Fisher Scientific) (43). Briefly, longer (34-37nt) oligos featuring the T7 promoter (17nt+1 G nt), the sgRNA designs targeting the promoter and the SBS region of SphK1, and a portion of the crRNA/tracrRNA constant region (15nt) were synthesized together as a template for the *in vitro* transcription **(Fig S2 (i)).** The following IVT (F1 and R1) oligos were designed as per the manufacturer instructions/kit manual **(Table S2).**

The initial PCR reaction was assembled as per the manufacturer instructions/kit manual for the generation of dsDNA templates of sgRNAs targeting either the HPRT gene (Control gRNA F1 and R1 Oligo Mix provided with the kit), the SphK1 Promoter (ProF1 and ProR1) or the SphK1 SBS region **(Table S3)**. The two-step PCR program was set for the above reaction: Initial denaturation at 98°C for 10 seconds (1X), Denaturation at 98°C for 5 seconds (32X) and Annealing at 55°C for 15 seconds (32X), Final extension at 72°C for 5 minutes (1X) and Hold at 4°C (1X).

The successive IVT reactions were setup with the PCR reaction mixes as follows **(Table S4)**. The reaction tubes were incubated in a water bath at 37°C for 120 minutes. 1μL of DNase I (at 1 U/µL) was added to each of the reaction mix immediately after the IVT reaction and both the reactions were incubated at 37°C for another 15 minutes. 0.5μL of the IVT reaction samples (diluted in 9.5μL Rnase Free Water) were analysed on a 2% agarose gel using specific protocols for separating RNA on agarose gels: Each of the samples were diluted 1:1 with (10μL of) a specific 2X RNA Gel Loading Dye (Thermo Fisher Scientific, Cat. No. R0641). All the sample mixes were heated/denatured at 70°C for 10 minutes and then chilled on ice till the samples were separated on a 2% agarose gel. The RNA content in each of the IVT reaction mix was directly measured using the Thermo Scientific™ NanoDrop™ 2000 Spectrophotometer.

### Screening the efficiency of the SphK1 sgRNAs using the Cas9 cleavage assay

Efficiencies of the sgRNAs designed to target the SphK1 gene were screened using the Guide-it sgRNA Screening Kit (Takara Bio USA, Inc., Cat. No. 632639) (44–47). An old genomic DNA stock dilution was utilized for the experiment. However, fresh genomic DNA could easily be extracted using the Extraction Buffers I and II provided with the kit. Around 1*10^6^ normal/untreated/control cells were seeded in a T-25 flask and allowed to grow till 80-90% confluency is reached (typically up to 36 hours). Genomic DNA from the confluent vials was harvested after incubation with the Extraction Buffers 1 and 2 provided in the same kit, according to the manufacturer’s instructions. The lysate was directly diluted 1:9 in Rnase Free Water (5μl of lysate + 45μl of Rnase Free Water). The genomic DNA content of the diluted lysate was directly measured using the Thermo Scientific™ NanoDrop™ 2000 Spectrophotometer. The respective target sequences of both the sgRNAs were first PCR amplified using the PCR reagents provided with the kit.

*In vitro* efficiencies of the sgRNAs designed to target the SphK1 gene were screened using the Guide-it sgRNA Screening Kit (Takara Bio USA, Inc., Cat. No. 632639). Entire volumes of all the PCR products were run on a 1% agarose DNA separation gel. The PCR products were eluted from the agarose DNA separation gel using the Geneall Expin^TM^ Gel SV elution kit (Free Sample; Catalog Number 102-110). The PCR amplified DNA content eluted from the agarose gel were directly measured using the Thermo Scientific™ NanoDrop™ 2000 Spectrophotometer. The following reaction was set up in parts (A and B) to monitor the Cas9 cleavage efficiency with the SphK1 sgRNA designs:

A. The target specific sgRNAs (approx. 90ng/μl; made after diluting 180ng/μl sgRNA dilutions in NFW in a 1:1 ratio) were incubated at 37°C for 5 minutes and then at 4°C (on ice) with the recombinant Cas9 nuclease provided with the kit in the following reaction **(Table S5).**
B. Cas9 cleavage assay was performed by assembling the following reaction **(Table S6)**.

All the reaction mixes were incubated in a Thermocycler using the following program set up, 37°C for 1 hr., 80°C for 5 min and 4°C Hold/Forever. The reactions were analyzed on a 1% agarose separating gel using a self-prepared DNA loading dye.

### Transfection of sgRNA plasmids in THP-1 cells using Electroporation

Macrophage-like THP-1 cells in the exponential phase of growth were plated in a 25cm^2^ flask at a density of 1 × 10^6^ cells/flask and were cultured overnight. Log-phased cells (4 × 10^6^) were split 24h before electroporation to ensure the cells are growing and healthy. Before electroporation, the cells were centrifuged and then resuspended in serum-and antibiotic-free RPMI-1640 and mixed with sgRNA plasmid DNA. The electroporated cells were removed from the dead cells and transferred into prewarmed complete medium. Electroporation was performed as described previously (48) with a Bio-Rad Gene 280 Pulser device. The cells were plated in 25cm^2^ flask and allowed to recover overnight. After overnight recovery, the media was removed and replenished with a fresh media, at which time the cells were ready for experimental studies.

### Infectivity assay

THP-1 cells (1 × 10^6^ cells/well), treated with 50 ng/ml of PMA (Sigma-Aldrich, USA) were seeded on glass coverslips in 6-well plates for 48h. They were infected and simultaneously treated with PF-543 (17µM) and or ABC294640 (70µM). After 48h, cells were Giemsa-stained, and intracellular parasite load was quantified microscopically by counting 10 random fields per coverslip as described previously by Pawar et al (49).

### qRT-PCR for gene expression analysis

Total RNA from infected macrophages was isolated using the TRIZOL reagent (Sigma-Aldrich, USA) and its concentration was determined by Nanodrop (Thermo Fischer, USA). cDNA was prepared from one microgram of RNase-free DNase treated total RNA using first-strand cDNA Synthesis Kit (Thermo Fischer Scientific, USA), as per manufacturer’s instructions, using random hexamer primers. The resulting cDNA was analyzed by quantitative real-time (qRT-PCR) RT-PCR (Applied Biosystems, 7500 Fast Real-Time PCR System, CA, USA) with gene-specific primers using PowerUp SYBR Green PCR Master Mix (Thermo Fisher Scientific, USA). The Primer sequences for IL-10, TNF-α, RNU6AP, JW, SphK1, and SphK2 are given in **Table S7**. Thermal profile for the real-time PCR was amplification at 50°C for 2 min followed by 40 cycles at 95°C for 15 sec, 60°C for 1 min and 72°C for 20 sec. Melting curves were generated along with the mean C_T_ values and confirmed the generation of a specific PCR product. Amplification of RNU6AP (RNA, U6 small nuclear 1; THP-1 cells) was used as internal control for normalization. The results were expressed as fold change of control (Uninfected samples (RNU6AP) using the 2^-ΔΔ*CT*^method. All samples were run in triplicates, including a no-template (negative) control for all primers used.

### Western blotting

Western blot analysis was done as described previously by Darlyuk et al., 2009 (50). Briefly, protein was isolated from the THP-1 cells by resuspending cell lysates in RIPA buffer. Before lysis, adherent macrophages were placed on ice and washed with PBS. Macrophages were scraped in the presence of RIPA lysis buffer containing 1% NP-40, 50mM Tris-HCl (pH 7.5), 150mM NaCl, 1mM EDTA (pH 8), 10mM 1,10-phenanthroline and phosphatase and protease inhibitors (Roche). After incubation, lysates were centrifuged for 15 min to remove insoluble matter.The proteins in the lysates were quantified, 80μg of the lysate was boiled (95¼C) for 5 min in SDS sample buffer and was subjected to electrophoresis on a 10% SDS-polyacrylamide gel. Proteins were then transferred onto nitrocellulose (NC) membrane using an electrophoretic transfer cell (Bio-Rad Laboratories, USA) at RT. The membrane was washed with 1× TBST solution three times and blocked with 5% BSA for 2h at RT. The blocked membrane was washed in TBST solution three times. The membrane was then incubated with primary monoclonal antibodies for Atg5, Beclin1, LC-3I/II, cytochrome c, Pro-caspase 3 and Caspase 9 in PBS-Tween 20 containing 5% BSA and incubated overnight at 4^ο^C. The blots were subsequently incubated with the secondary antibody conjugated to horseradish peroxidase at 1:3000 dilution in 5% PBS-Tween 20 for 2h at RT. Enhanced chemiluminescence reaction was used for the detection of the blot. The results were expressed as fold change and quantitated by using AlphaEaseFC image analysis software (Alpha Innotech). The data were expressed as mean ± SD of three independent experiments, and the representative image of one experiment is shown.

### Immunofluoroscence

THP-1 macrophages plated on coverslips were infected with *Leishmania donovani* for 48h. After blocking with 3% BSA, cells were stained using anti-SphK1 (NBP1-69235, Novus) and anti-SphK2 (D885, Cell Signalling) antibodies. Slides were washed twice with PBS-T and once with PBS, and probed with Alexa Fluor 488 (green)-conjugated goat anti-mouse IgG antibodies (1:200; Molecular Probes) at RT for 1h. Following washing, the slides were mounted with Hoechst stain and visualized under a confocal laser scanning microscope (Olympus FluoViewTM FV1000 with objective lenses PLAPON ×60 O, NA-1.42) at an excitation wavelength of 556 nm. Images were processed via NIS-Elements software version 4.50. The mean fluorescence intensities were plotted using GraphPad Prism version 8.0. The experiment was performed thrice.

### NBD-Sph assay

A fluorescent molecule: omega (7-nitro-2–1, 3-benzoxadiazol-4-yl [2S,3R,4E]-2-amino octadec-4-ene-1,3-diol [NBD–Sphingosine (NBD-Sph); Avanti Polar Lipids]) was used as a substrate, and the effect of SphK mutants (THP-1 transfected with SphK sgRNA plasmids) on the conversion of NBD-Sph to NBD-S1P by the infected macrophages was evaluated. Toward this, *Leishmania-*infected macrophages (20:1 MOI) were transfected with SphK Crispr/Cas9 mutants, for 72h. A total of 100μL (1 × 108 cells) of this culture was washed with incomplete RPMI (iRPMI), and incubated with 10μM NBD-Sph, at 37°C for 60 min. After incubation, the cells were lysed with RIPA lysis buffer to obtain host-free parasites. The infected macrophages were thoroughly mixed with 260μL of methanol and 400μL of chloroform: methanol (1:1), followed by adding 16μL of 7 M NH4OH, 400μL of chloroform, and 300μL of 1.5 M KCl. The lipids were separated by centrifugation at 17,000 × g for 5 min. A 100μL aliquot of the upper (aqueous) phase was transferred to a black 96-well flat-bottom plate (Corning). Fluorescence intensity of the aqueous phase containing NBD-S1P, in SphK mutant transfected and un-transfected samples, was measured at an excitation/emission wavelength of 485 nm/530 nm using Varioskan LUX Multimode Microplate Reader (ThermoFisher Scientific). The data obtained from 3 independent experiments were plotted using GraphPad Prism version 8.0 software. The fluorescence intensity of NBD-S1P was measured as described above.

### Statistical analysis

Fold-expression (qRT-PCR and densitometric analysis) and intracellular parasite burden were represented as mean ±SD. Each experiment was repeated three times in separate sets. Statistical differences were determined using Student’s unpaired 2-tailed *t*-test. All statistics were performed using GraphPad Prism Version 5.0 (GraphPad Software, USA). p ≤ 0.05 was considered significant [* (P<0.01 to 0.05), ** (P< 0.001), *** (P< 0.0001), ns (P≥ 0.05)].

## Acknowledgments

Singh S conceptualized the work, designed the experiments, interpreted the data and edited the manuscript and acquired the funding to support the work. EM executed the experiments, performed the data interpretation and wrote the first draft of the manuscript. EM, JS, SM, AG, SP and WD performed immunological studies, microscopy, molecular assays, mammalian culture, sgRNA cloning and transfection. This study was supported by CSIR Scientist Pool Officer (SRA) funding agency research grant (13(9160-A)2021-Pool) provided to Dr. Evanka Madan. JS was supported by BioCARe Women Scientist. SS is a recipient of the National Bio Scientist Award. All authors contributed to the article and approved the submitted version. The work in Shailja Singh’s lab at Special Centre for Molecular Medicine (SCMM), JNU, is supported by IRHPA IPA/2020/000007, Department of Science and Technology and Drug and Pharmaceuticals Research Program (DPRP) (Project No. P/569/2016-1/TDT, SS). SM and WD acknowledge Shiv Nadar University for access to their instrumentation facilities and thank the university for research support extended to them through internal foundation funding. We would like to acknowledge Prof. Md. Imtiyaz Hassan (Professor, Structural Biology Laboratory, Jamia Millia Islamia) for providing us with the Human SphK1 clone using which we raised Human SphK1 Antibody in Swiss mice (obtained from the Central Laboratory Animal Resources, Jawaharlal Nehru University, New Delhi) using standard protocol approved by IAEC, JNU). The funders had no role in the study design; collection, analysis, and interpretation of data; in the writing of the report; and in the decision to submit this article for publication.

## Plagiarism Check

This manuscript has been screened for plagiarism and found to be original.

## References

1. Anand PK, Kaul D, Grover S, Vanakkam A, Natarajan K, Kaul A, Saini A, Agrewala JN. (2014). Science signaling. Sci Signal. 7(331):ra45.

2. Pucadyil TJ, Chattopadhyay A. (2006). Biochimica et Biophysica Acta – Biomembranes. Biochim Biophys Acta. 1758(2):161–171.

3. Blaho VA, Hla T. (2014). Sphingosine 1-phosphate signaling in immunity. J Clin Invest. 124(6):2277–2283.

4. Spiegel S, Milstien S. (2011). The outs and the ins of sphingosine-1-phosphate in immunity. Nat Rev Mol Cell Biol. 12(9):569–579.

5. Rivera J, Proia RL, Olivera A. (2008). The S1P–S1P1 axis in immunity. J Immunol. 181(9):6664–6669.

6 . Matloubian M, Lo CG, Cinamon G, Lesneski MJ, Xu Y, Brinkmann V, Allende ML, Proia RL. (2004). Lymphocyte egress and S1P1 receptor. Nature. 427:355–360.

7. Oskeritzian CA, Price MM, Hait NC, Milstien S, Pyne S, Pyne NJ. (2007). FTY720 inhibits mast cell functions. FASEB J. 21(14):3256–3268.

8. Price MM, Goodarzi K, Khan MF, Nguyen L, Candia J, Deepe GS Jr. (2013). The S1P pathway in fungal allergy. J Allergy Clin Immunol. 131(3):808–815.

9. Maceyka M, Sankala H, Hait NC, Le Stunff H, Liu H, Toman R, Collard EJ, Thangada S, Spiegel S. (2012). S1P and lipid metabolic signalling. Biochim Biophys Acta. 1818(9):2388–2397.

10. Bhardwaj S, Banerjee S, Baldi A, Chadha V, Kumar A, Kumar A, Tripathi BK. (2022). Sphingosine kinase and immune responses in infection. Front Immunol. 13:845394.

11. Cáceres M, Alonso A, Goni FM. (2020). Sphingosine-1-phosphate in vascular biology. Cells. 9(5):1245.

12. Ran FA, Hsu PD, Wright J, Agarwala V, Scott DA, Zhang F. (2013). Genome engineering using the CRISPR–Cas9 system. Nat Protoc. 8(11):2281–2308. doi:10.1038/nprot.2013.143

13. Sanjana NE, Shalem O, Zhang F. (2014). Improved vectors and genome-wide libraries for CRISPR screening. Nat Methods. 11:783–784.

14. Cong L, Ran FA, Cox D, Lin S, Barretto R, Habib N, Hsu PD, Wu X, Jiang W, Marraffini LA, Zhang F. (2013). Multiplex genome engineering using CRISPR/Cas systems. Science. 339(6121):819–823. doi:10.1126/science.1231143

15. Wattenberg BW, Sabbadini RA. (2003). Roles of sphingosine-1-phosphate in cancer and angiogenesis. Biochim Biophys Acta. 1585:112–117.

16. Maceyka M, Payne SG, Milstien S, Spiegel S. (2005). Sphingosine kinases in immunity and cancer. Adv Cancer Res. 95:1–51.

17. Billich A, Bornancin F, Dévay P, Mechtcheriakova D, Urtz N, Baumruker T. (2003). Phosphorylation of the sphingosine analogue A-kinase anchoring protein. FEBS Lett. 554(1–2):32–38.

18. Tousif S, Bolla PK, Thaisrivongs S, Chandrani M, Moideen K, Ahmed M, Ranganathan V, Sultana R, Ahmed A, Ahmad I, Reddy MC, Das G. (2017). Nanoparticle-formulated curcumin prevents posttherapeutic reactivation and reinfection with *Mycobacterium tuberculosis* following isoniazid therapy. Front Immunol. 8:420. doi:10.3389/fimmu.2017.00420

19. Sundar S, Chakravarty J. (2018). Current treatment of visceral leishmaniasis. Expert Opin Pharmacother. 19(3):261–267.

20. Ghosh K, Mandal G, Gore K. (2013). Cytokine regulation in *Leishmania* pathogenesis—a review. Immunobiology. 218(4):551–557.

21. Gupta G, Kumar R, Dey R, Tiwary P, Natarajan G, Salotra P. (2020). Inflammatory response modulation by sphingolipids in parasitic infections. Front Cell Infect Microbiol. 10:80. doi:10.3389/fcimb.2020.00080

22. Maceyka M, Spiegel S. (2014). Sphingolipid metabolites in inflammatory disease. Nature. 510(7503):58–67.

23. Shen Y, Zhang W, Gao X, Huang Q. (2014). Sphingosine kinase 1 knockdown triggers apoptosis in macrophages. Cell Death Dis. 5(5):e1025.

24. Nandan D, Reiner NE. (2005). Manipulation of macrophage signalling pathways by intracellular protozoan parasites. Trends Microbiol. 13(8):404–410.

25. Soliman M, Soliman E, Reslan H, Fathy G, Dkhil MA, Al-Quraishy S. (2020). Targeting host cell death pathways for therapy in parasitic infections. Parasite Immunol. 42(4):e12704. doi:10.1111/pim.12704

26. Hait NC, Allegood J, Maceyka M, Strub GM, Harikumar KB, Singh SK, Luo C, Muthusamy BP, Hla T, Kordula T, Milstien S, Spiegel S. (2014). Sphingosine-1-phosphate in the nucleus regulates histone acetylation. Science. 344(6185):1244–1248. doi:10.1126/science.1250140

27. Schnute ME, McReynolds MD, Kasten TJ, Younts TJ, Yadegari H, Radek JT, Rupprecht E, Dumaual N, Oskeritzian CA, Price MM, Hait NC, Jiang B, Milstien S, Pyne S, Pyne NJ, Edsall LC, Guerard EJ, Lacapra S, Johnson KM, Ramanathan R, et al. (2012). Modulation of SphK1/2 using isoform-selective inhibitors. J Pharmacol Exp Ther. 343(2):474–488. doi:10.1124/jpet.112.197921

28. Ohta H, Ishizuka T, Hamano H, Doi Y, Sato K, Nakamura K, Sugiyama T, Takabayashi K. (2004). Sphingosine-1-phosphate regulates chemotaxis and leukocyte recruitment. J Biol Chem. 279(20):20414–20424. doi:10.1074/jbc.M310048200

29. Yogesh R, Horn D, Rahman A. (2021). Sphingolipids in protozoan infection: Therapeutic targets. Trends Parasitol. 37(2):127–142. doi:10.1016/j.pt.2020.11.012

30. Lima-Junior DS, Costa DL, Carregaro V, Cunha LD, Silva AL, Mineo TW, Alves LS, Rosa J, Martin CJ, de Freitas MS, Feminino JS, Palma C, Rangel TP, Zaccone P, Mansour E, Oliveira SC, Lima-Junior DS. (2011). Autophagy and parasite clearance. Cell Host Microbe. 10(5):471–483. doi:10.1016/j.chom.2011.09.003

31. Spiegel S, Milstien S. (2011). The outs and the ins of sphingosine-1-phosphate in immunity. Nat Rev Immunol. 11(5):403–415. doi:10.1038/nri2974

32. Bektas M, Spiegel S. (2004). Sphingosine-1-phosphate and apoptosis. Biochim Biophys Acta. 1585(2-3):193–201. doi:10.1016/j.bbalip.2003.09.003

33. Ghosh S, Subramanian S, Rahman A, Ghosh A, Biswas N, Sarkar A, Fernandopulle P, Rathod PK, Bandyopadhyay S. (2012). Sphingosine kinase 1 regulates the macrophage inflammatory response. J Immunol. 188(1):124–133. doi:10.4049/jimmunol.1101629

34. Wang F, Cao R, Fan C, Xu G, Teng L, Cao W, Qi X, Wang R, Xu H, Liang C, Ruan C. (2014). S1P signaling drives Th2 polarization. Nat Commun. 5:3566. doi:10.1038/ncomms4566

35. Estrada LD, Moras M, Jiang H, Chapman NM, Belt R, Bricker TL, Patel N, Urban JF Jr, Hla T, Pearce EJ. (2012). Sphingosine-1-phosphate promotes Th2 inflammation by inhibiting IFN-γ signaling. Nat Immunol. 13:1256–1264. doi:10.1038/ni.2468

36. Hait NC, Maiti A. (2017). The role of sphingosine-1-phosphate and ceramide-1-phosphate in inflammation and cancer. Mediators Inflamm. 2017:4806541. doi:10.1155/2017/4806541

37. Snider AJ, Kawamori T, Bradshaw SG, Orr G, Alekseev A, Kohama T, Khan O, Simpson RJ, Wang Z, Milstien S, Spiegel S. (2008). Sphingosine kinase: Role in regulation of bioactive sphingolipid mediators in inflammation. Biochimie. 90(4):596–612. doi:10.1016/j.biochi.2007.10.006

38. Pyne NJ, Pyne S. (2011). Sphingosine 1-phosphate and cancer. Nat Rev Cancer. 10(7):489–503. doi:10.1038/nrc3088

39. Liu H, Toman RE, Goparaju SK, Maceyka M, Nava VE, Sankala H, Mitsutake S, Spiegel S. (2000). Sphingosine kinase type 2 is a putative BH3-only protein that induces apoptosis. J Biol Chem. 275(11):7601–7604. doi:10.1074/jbc.275.11.7601

40. Sundar S, Chakravarty J. (2015). An update on pharmacotherapy for leishmaniasis. Expert Opin Pharmacother. 16(2):237–252. doi:10.1517/14656566.2015.986729

41. Ponte-Sucre A, Gamarro F, Dujardin J-C, Barrett MP, López-Vélez R, García-Hernández R, Puerta C, Pountain A, Uliana S, Morales EK, Hill L, Sundar S, von Schrével J, Maes L, Bañuls AL. (2017). Drug resistance and treatment failure in leishmaniasis: A 21st century challenge. PLoS Negl Trop Dis. 11(12):e0006052. doi:10.1371/journal.pntd.0006052

42. Hengst JA, Dick TE, Smith CD, Yun JK. 2020. Analysis of selective target engagement by small-molecule sphingosine kinase inhibitors using the cellular thermal shift assay (CETSA).Cancer Biol Ther 21:841 852. 10.1080/15384047.2020.1798696.

43. Takara Bio USA, Inc. (2016). Guide-it Precision gRNA Synthesis Kit User Manual: For the generation of full-length guide RNA (gRNA) for use with CRISPR/Cas9-mediated genome editing. Takara Bio USA, Inc., Mountain View, CA.

44. Anders C, Jinek M. (2014). In vitro enzymology of Cas9. Methods Enzymol. 546:1–20. doi:10.1016/B978-0-12-801185-0.00001-5

45. Zhang K, Deng R, Li Y, Zhang L, Li J. (2016). Cas9 cleavage assay for pre-screening of sgRNAs using nicking triggered isothermal amplification. Chem Sci. 7(8):4951– 4957. doi:10.1039/C6SC01355D

46. Karmakar S, Behera D, Baig MJ, Molla KA. (2021). In vitro Cas9 cleavage assay to check guide RNA efficiency. In: Editor(s)], Methods in Molecular Biology, Vol 2384. Springer; p. 23–39. doi:10.1007/978-1-0716-1657-4_3

47. Takara Bio USA, Inc. 2018. Guide-it™ sgRNA In Vitro Transcription and Screening Systems User Manual. Takara Bio USA, Inc., Mountain View, CA.

48. Kapler GM, Coburn CM, Beverley SM. (1990). Stable transfection of the human parasite *Leishmania major* delineates a 30-kilobase region sufficient for extrachromosomal replication and expression. Mol Cell Biol. 10(3):1084–1094.

49. Pawar H, Puri M, Fischer-Weinberger R, Madhubala R, Zilberstein D. (2019). The arginine sensing and transport binding sites are distinct in the human pathogen *Leishmania*. PLoS Negl Trop Dis. 13(1):e0007304. doi:10.1371/journal.pntd.0007304

50. Darlyuk I, Goldman A, Roberts SC, Ullman B, Rentsch D, Zilberstein D. (2009). Arginine homeostasis and transport in the human pathogen *Leishmania donovani*. J Biol Chem. 284(30):19800–19807. doi:10.1074/jbc.M109.007245

