## Supplementary figures and images for "CRISPR/Cas9-mediated identification of human macrophage SphK1 as druggable target for development of anti-leishmanial chemotherapeutics"

### Supplementary Figure 1

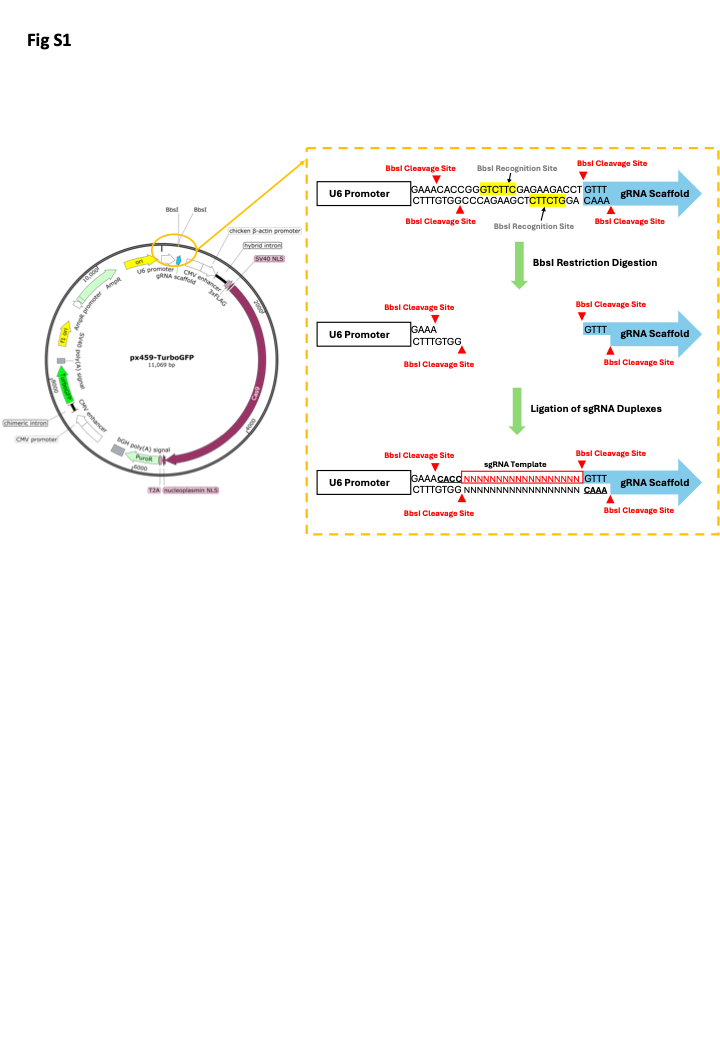

### Supplementary Figure 2

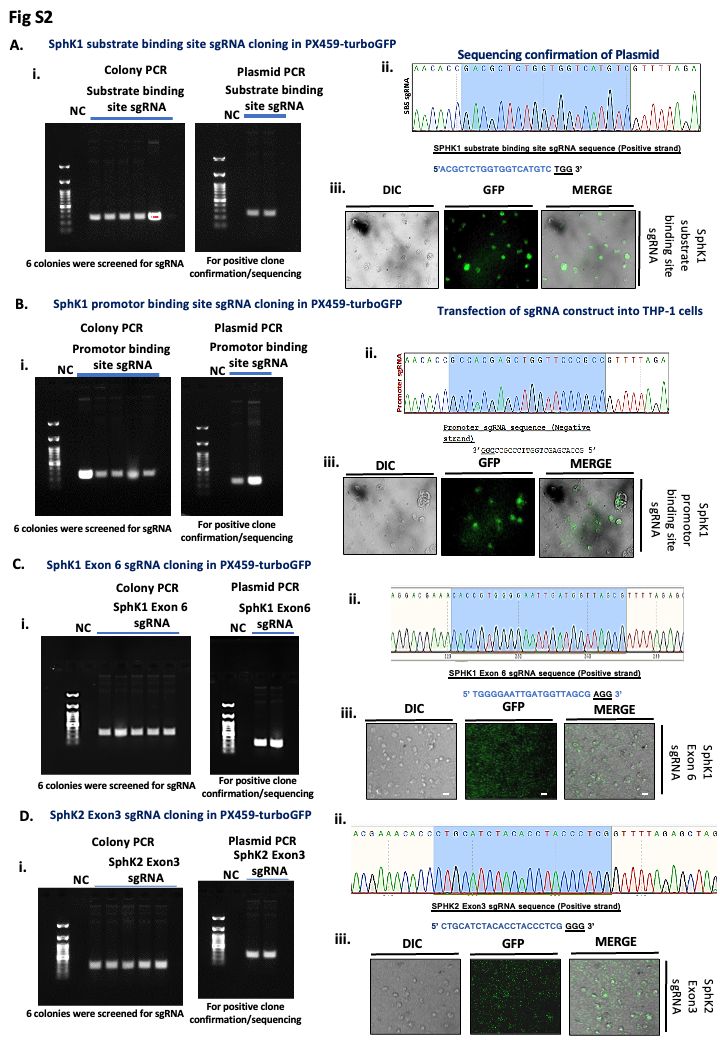

### Supplementary Figure 3

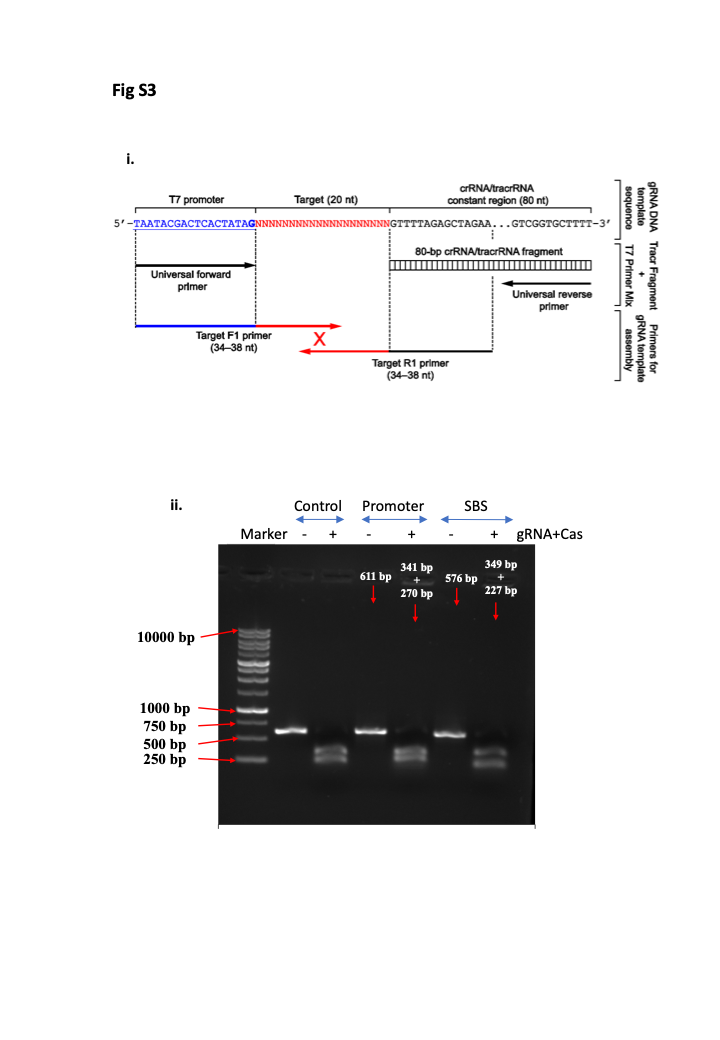
