## Supplementary Tables and Legends for "CRISPR/Cas9-mediated identification of human macrophage SphK1 as druggable target for development of anti-leishmanial chemotherapeutics"

**Supplementary Figure Legends:**

**Fig S1. Cloning Strategy for SphK1 sgRNAs targeting the Substrate-Binding Site and Promoter Region**

**(i)** Schematic representation of the PX459 vector used for cloning SphK1-specific sgRNAs. **(ii)** Cloning of sgRNAs targeting the SphK1 substrate-binding site (SBS) and promoter region into the PX459 vector backbone.

**Fig S2. Validation of SphK1 and SphK2 sgRNA constructs and transfection efficiency in THP-1 macrophages**

sgRNAs targeting the substrate-binding site, promoter, and exon 6 regions *of* SphK1, and exon 3 of SphK2*,* were cloned into the PX459-TurboGFP vector. **(i)** Colony PCR using U6 forward and sgRNA-specific reverse primers confirmed the presence of correct inserts (~222 bp). **(ii)** Sanger sequencing validated the fidelity and orientation of the cloned sgRNAs. **(iii)** GFP fluorescence observed in THP-1 macrophages 48–72 h post-transfection confirmed successful delivery and expression of the CRISPR-Cas9 constructs.

**Fig S3. Evaluation of SphK1 sgRNA efficiency using *in vitro* Cleavage Assay**

**(i)** PCR amplification of double-stranded sgRNA templates using synthetic forward (F1) and reverse (R1) oligonucleotides in combination with the Tracr fragment and T7 primer mix.
**(ii)** *In vitro* cleavage assay performed using ~100ng of either the kit-provided control DNA template or DNA templates corresponding to the promoter and substrate-binding site (SBS) regions of the SphK1 gene. Each reaction included ~100ng of *in vitro* transcribed sgRNA and ~250ng of recombinant Cas9 protein, incubated under optimal conditions to assess sgRNA-mediated cleavage efficiency.

**Supplementary Tables & Table Legends:**

**Table S1: Forward and Reverse sequences of the best sgRNA designs targeting the promoter region, SBS region and exon-6 of the hSphK1 gene along with exon-3 of the hSphK2 gene**

| sgRNA ID | Forward Sequence | Reverse Sequence |
| --- | --- | --- |
| SphK1 pro-sgR | 5’-GCCACGAGCTGGTTCCCGCC-3’ | 5’-GGCGGGAACCAGCTCGTGGC-3’ |
| SphK1-SBS-sgR | 5’-GACGCTCTGGTGGTCATGTC-3’ | 5’-GACATGACCACCAGAGCGTC-3’ |
| SphK1-Exon-6-sgR | 5’-GTGGGGAATTGATGGTTAGCG-3’ | 5’-AAACCGCTAACCATCAATTCC-3’ |
| SphK2-Exon-3-sgR | 5’-CTGCATCTACACCTACCCTCG-3’ | 5’- CGAGGGTAGGTGTAGATGCAG-3’ |

**Table S2: The IVT (*In vitro transcription)*** **(F1 and R1) oligos sequences and their respective lengths**

| **Oligo ID** | **Oligo Sequence (5’-3’)** | **Oligo Length (bp)** |
| --- | --- | --- |
| SphK1proF1_geneart | TAATACGACTCACTATAGCCACGAGCTGGTTCCCG | 35 |
| SphK1proR1_geneart | TTCTAGCTCTAAAACGGCGGGAACCAGCTCGTGG | 34 |
| SphK1SBSF1_geneart | TAATACGACTCACTATAGGACGCTCTGGTGGTCATGT | 37 |
| SphK1SBSR1_geneart | TTCTAGCTCTAAAACGACATGACCACCAGAGCGT | 34 |

**Table S3: Conditions for the** **initial PCR reaction for the generation of dsDNA templates of sgRNAs targeting either the HPRT gene (Control gRNA F1 and R1 Oligo Mix provided with the kit), the SphK1 Promoter (ProF1 and ProR1) or the SphK1 SBS region**

| **Component** | **Volume for Control (µl)** | **Volume for Promoter (µl)** | **Volume for**  **SBS (µl)** |
| --- | --- | --- | --- |
| Phusion™ High-Fidelity PCR Master Mix | 12.5 | 12.5 | 12.5 |
| Tracr Fragment+T7 Primer Mix | 1 | 1 | 1 |
| 0.3µM Target F1-R1 oligonucleotide mix^*#^ | 1 | 1 | 1 |
| Nuclease-free water | 10.5 | 10.5 | 10.5 |

**Table S4: PCR reaction conditions for successive IVT reactions**

|  | **Volume for sgPromoter (µl)** | **Volume for sgSBS (µl)** |
| --- | --- | --- |
| NTP mix  (25 mM each of ATP, GTP, CTP, UTP in Tris buffer) | 16 | 16 |
| gRNA DNA template  (from PCR assembly) | 12 | 12 |
| 5X TranscriptAid™ Reaction Buffer | 8 | 8 |
| TranscriptAid™ Enzyme Mix | 4 | 4 |

**Table S5: The following reaction conditions for target specific sgRNAs**

| **Reagent** | **Volume (μl)** |
| --- | --- |
| Target-specific sgRNA (90 ng/μl) or Control sgRNA (50 ng/μl) | 1 |
| Guide-it Recombinant Cas9 Nuclease (500 ng/μl) | 0.5 |
| Total | 1.5 |

**Table S6: The following reaction conditions for Cas9 cleavage assay**

| **Reagent** | **Volume with Control Fragment (20ng/μl)** | **Volume with sgR_Pro (app. 90 ng/μl) and Promoter Fragment**  **(111ng/μl)** | **Volume for sgR_SBS (app. 90 ng/μl) and SBS Fragment (112ng/μl)** |
| --- | --- | --- | --- |
| PCR reaction solution | 5 | 1 | 1 |
| 15X Cas9 Reaction Buffer | 1 | 1 | 1 |
| 15X BSA | 1 | 1 | 1 |
| RNase Free Water | 6.5 | 10.5 | 10.5 |
| Cas9/sgRNA mix (from Step A above) | 1.5 | 1.5 | 1.5 |
| Total | 15 | 15 | 15 |

**Table S7: Primer sequences of target genes mentioned along with their annealing temperatures (Tm)**

| **Primer** | **Sequence (5’-3’)** | **Annealing Temp** |
| --- | --- | --- |
| IL-10 | Forward 5′–TTAAGGGTTACCTGGGTTGC-3′  Reverse 5′-TGAGGGTCTTCAGGTTCTCC-3′ | 60^o^C |
| TNF-α | Forward 5′-CCTCTCTCTAATCAGCCCTCTG-3′  Reverse 5′-GAGGACCTGGGAGTAGATGAG-3′ | 60^o^C |
| RNU6AP | Forward 5′-GGCCCAGCA GTACCTGTTTA-3′  Reverse 5′-AGATGGCGGAGGTGCAG-3′ | 60^o^C |
| JW | Forward 5′-CCTATTTTACACCAACCCCCAGT-3′  Reverse 5′-GGGTAGGGGCGTTC TGCGAAA-3′ | 60^o^C |
| SphK1 | Forward 5’-GGAGACCGCCATCCAGAA-3’  Reverse 5’-CTCATAGCCAGCATAATGGTTCAA-3’ | 59^o^C |
| SphK2 | Forward 5′–CCCCGGTTGCTTCTATTGGT-3′  Reverse 5′-CCCACAGGCAGGCTGATTTA-3′ | 60^o^C |
